# Cultural Selection Shapes Network Structure

**DOI:** 10.1101/464883

**Authors:** Marco Smolla, Erol Akçay

**Affiliations:** Department of Biology, University of Pennsylvania, Philadelphia, PA, 19104, USA

## Abstract

Cultural evolution relies on the social transmission of cultural traits across a population, along the ties of an underlying social network that emerges from non-random interactions among individuals. Research indicates that the structure of those interaction networks affects information spread, and thus a population’s ability for cumulative culture. However, how network structure itself is driven by population-culture co-evolution remains largely unclear. We use a simple but realistic model of complex dynamic social networks to investigate how populations negotiate the trade-off between acquiring new skills and getting better at existing skills, and how this trade-off, in turn, shapes the social structure of the population. Our results reveal unexpected eco-evolutionary feedback from culture onto social network structure and vice versa. We show that selecting for generalists (favouring a broad repertoire of skills) results in sparsely connected networks with highly diverse skill sets, whereas selecting for specialists (favouring skill proficiency) results in densely connected networks and a population that specializes on the same few skills on which everyone is an expert. Surprisingly, cultural selection for specialisation can act as an “ecological trap” where it can take a long time for a specialist population to adapt to a generalist world. Our model advances our understanding of the complex feedbacks in cultural evolution and demonstrates how individual-level behaviour can lead to the emergence of population-level structure.

## Introduction

Cultural evolution relies on the transmission of cultural traits, such as skills, technologies, believes, and ideas among individuals of a population and as such it is fundamentally social^1–3^. Therefore, the amount and nature of culture that populations accumulate is expected to depend on the structure of social groups culture evolves in. Theoretical and empirical studies hypothesized and demonstrated a direct link between group demography and culture showing, for example, that a population’s ability to accumulate cultural traits depends on its size, mobility, and density^4–7^. In experiments, larger groups maintained higher cultural complexity than smaller groups^8^, and when faced with a complex task, groups accumulated beneficial solutions more readily than single individuals^9;10^.

In this paper, we focus on the reciprocal feedbacks between the fine-scale social structure of a population and its cumulative culture. As information is transmitted between interacting individuals, its flow follows the ties of a social network^11–14^. Social interactions are often non-random, mediated by diverse factors and processes such as spatial heterogeneity^15^, homophily (e.g. behaviour matching)^16^, or social inheritance^17^, all of which lead to non-random opportunities for information to spread. The processes that structure social networks can thus affect how culture is transmitted and accumulated in populations, and therefore the adaptive properties of cumulative culture.

In well connected populations (high degree, short path length) information can spread faster and farther^7;18;19^, a prediction that is borne out in humans^10;20^ and non-human species^21–24^. Such networks might be optimal where group activity requires coordination, for example, in regards to social norms^25^ and rituals^26^. Faster information transmission, however, is not universally adaptive for groups. Fast convergence on a single solution can short circuit a thorough exploration of the solution space, and cause a group to settle for suboptimal solutions^27;28^. On the other hand, networks with slower information diffusion can increase the chance to collectively discover a global maximum^27;29^, innovate a wider variety of solutions to a given problem^27;30^, or retain more information in a collective memory-retrieval situation^31^. The desirable network structure (from the group’s perspective) thus depends on the collective problem to be solved, or the required diversity of solutions — a trade-off between cultural convergence and cultural diversity, broad exploration versus swift coordination^28;30;32^.

While the effect of network structure on the dynamics of cultural traits is well-studied, the converse effect, the effect of cultural dynamics on network structure remains unexplored. The above discussion implies that populations facing ecological pressures that require different types of cultural adaptations might be expected to evolve different network structures. In particular, if traits that affect how individuals form connections in their network are heritable (genetically or culturally), network structure can evolve under natural or cultural selection. The evolving network structure, in turn, will determine the nature of cumulative culture in the population, which can further affect the future evolution of network structure. We show in this paper that this “eco-evolutionary” feedback between cumulative culture and network structure can have unexpected consequences.

Our approach is based on a simple, yet generally applicable and realistic model of dynamic social networks^17^, where individuals make connections either by inheriting them from their parents (or other role models) or randomly. The probabilities of making these two types of connections determine the average degree, and whether the network is well-connected and has short path lengths, or is highly clustered with long path lengths. Previous work has shown that co-evolution of these linking traits with social behaviours can lead to unexpected dynamics such as the collapse of cooperation due to its effect on network evolution^33^. Here, we model the co-evolution of network structure with culture in a population where individuals socially acquire traits to cope with their environment. As social learning only occurs between connected individuals, an individual’s neighbourhood affects what it can learn, and so, how it will cope with the environment, intrinsically linking network and cultural dynamics. Given that network topology affects information flow, and thus what individuals can learn, we expect that different network topologies emerge in response to different requirements on cultural knowledge. To test this, we compare two different worlds: a *generalist world* that favours individuals with a broad selection of skills (e.g. societies where each individual contributes in similar ways to the subsistence of the group^26^), and a *specialist world* that favours individuals with high proficiency in at least one skill (e.g. societies with high division of labour that allows specialisation, which requires a lot of time engaging with only one trait^34–36^).

## Methods

### The model

We model populations of *N* asexually reproducing individuals, with overlapping generations, in a world with *T* learnable cultural traits. These traits relate to e.g. subsistence and social norms and, therefore, are relevant to an individual’s survival and reproduction. Traits are assumed to be equal in payoff and can be acquired independent from each other, as we are interested in how much and how well individuals learn but not what they learn.

Time is divided into rounds. Each round consists of three steps: (1) one randomly selected individual leaves the population, (2) a parent is selected and a new individual added to the population, and (3) the new individual acquires traits through innovation and copying. During learning the new individual has 100 alternating asocial and social learning attempts, allowing her to either acquire new traits or improve proficiency in those she already possesses. Restricting learning to a phase early in life is based on observations that children in hunter-gatherer societies acquire most skills prior to adolescence^26;37^. At birth, an individual’s proficiency *l* is zero for all traits (i.e. *l_t_* = 0, *t* ∈ *T*). Trait proficiency increases through successful learning. When a new trait is acquired through asocial (innovation) or social (copying) learning proficiency of trait *t* increases from *l_t_* = 0 to *l_t_* = 1. As the individual’s repertoire size *R_i_* is the number of non-zero trait proficiencies *l*, trait acquisition increases *R_i_*. To become better at performing a trait repeated engagement with it is required, as learning takes time^38–41^. Therefore, proficiency increases with each successful asocial or social learning attempt of the same trait so that 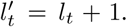 Because the number of learning turns is limited and attention to one trait limits attention to other traits, there is a trade-off between becoming good at a trait and learning many traits. Hence, trait proficiency and repertoire size are negatively related.

The probability for successful *asocial learning* of a trait*t* is

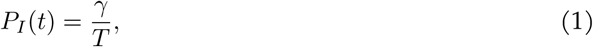

where *γ* is a fixed innovation success probability.

The probability for successful *social learning* of a trait *t* is

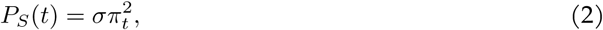

where *σ* is a fixed copying success probability, and *π_t_* is the probability that an individual observes a certain trait in its neighbourhood, which is

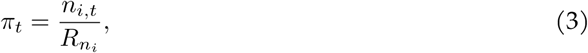

where *n_i,t_* is the number of *i’*s neighbours with trait *t*, and *R_n_i__* is the sum of repertoire sizes of *i*’s neighbours. Our assumption that an individual is not actively choosing a trait to learn is based on observations in traditional societies where children acquire knowledge through playful work^42^ or by helping their parents with subsistence tasks^37;41;43^. The traits they observe are those that are performed in their vicinity.

The use of a quadratic function in eq. 2 represents the need for more than a single observation of a trait for successful social learning. The rationale is that a trait needs to be sufficiently common in the neighbourhood to be observed but also sufficiently often performed so that the individual receives enough exposure to learn the trait. Eq. 2 relates to Simpson’s index^44^, a measure of diversity representing the probability that two randomly picked items of a population belong to the same type. Like Simpson’s index, *P_S_* behaves parabolic, where larger trait diversity is stronger punished than lower diversity. Therefore, *P_S_* (*t*) is highest where all neighbours perform only trait *t* (see electronic supplementary material S1). However, as trait diversity in *i’*s neighbourhood increases trait exposure decreases making it less likely to observe its performance sufficiently long to learn about it socially (Fig. S2).

As eq. 2 shows acquiring a trait socially is more likely if social learning is easy (large *σ*), trait *t* is common among neighbours (large *n_i,t_*), and if neighbours possess few traits (small *R_n_i__*). Furthermore, we assume that an individual cannot surpass the proficiency of the observed individuals, and thus *P_S_*(*t*) = 0 where all neighbours have proficiency equal to or less than that of the individual *i* for trait *t*.

Subsequent to the learning phase, we calculate an individual’s lifetime success score, or payoff *W*. Its magnitude depends on whether the individual acquired traits according to its environment. Individuals face one of two environments. In the *generalist world* individuals benefit from acquiring a variety of traits and so an individual’s payoff is equivalent to its repertoire size, *W_i_* = *R_i_*. In contrast, in the *specialist world* individuals benefit from becoming highly proficient in one trait. Here, an individual’s payoff is equivalent to the highest trait proficiency in its repertoire, *W_i_* = max(**L**_*i*_), where **L**_*i*_ is a vector of *i’*s proficiencies. The two environments can represent a variety of contexts, such as foraging. The *generalists* might forage on ephemeral, easy to handle but highly diverse resources, whereas the *specialists* might forage on stable, less diverse but hard to handle resources.

A new simulation round starts with the removal of a random individual. A survivor is selected as a parent to replace the individual relative to its payoff *W_i_*.

### Model iterations

#### Topology effects on culture

To establish a baseline for the effect of network topology on cultural dynamics in our model we begin with a set of *static*, *regular networks* (ring). We use different neighbourhood sizes (1,3, and 10) to alter topology (degree, clustering, average path length) and measure its effect on average repertoire size and average highest proficiency (Fig. 1).

**Figure 1:**
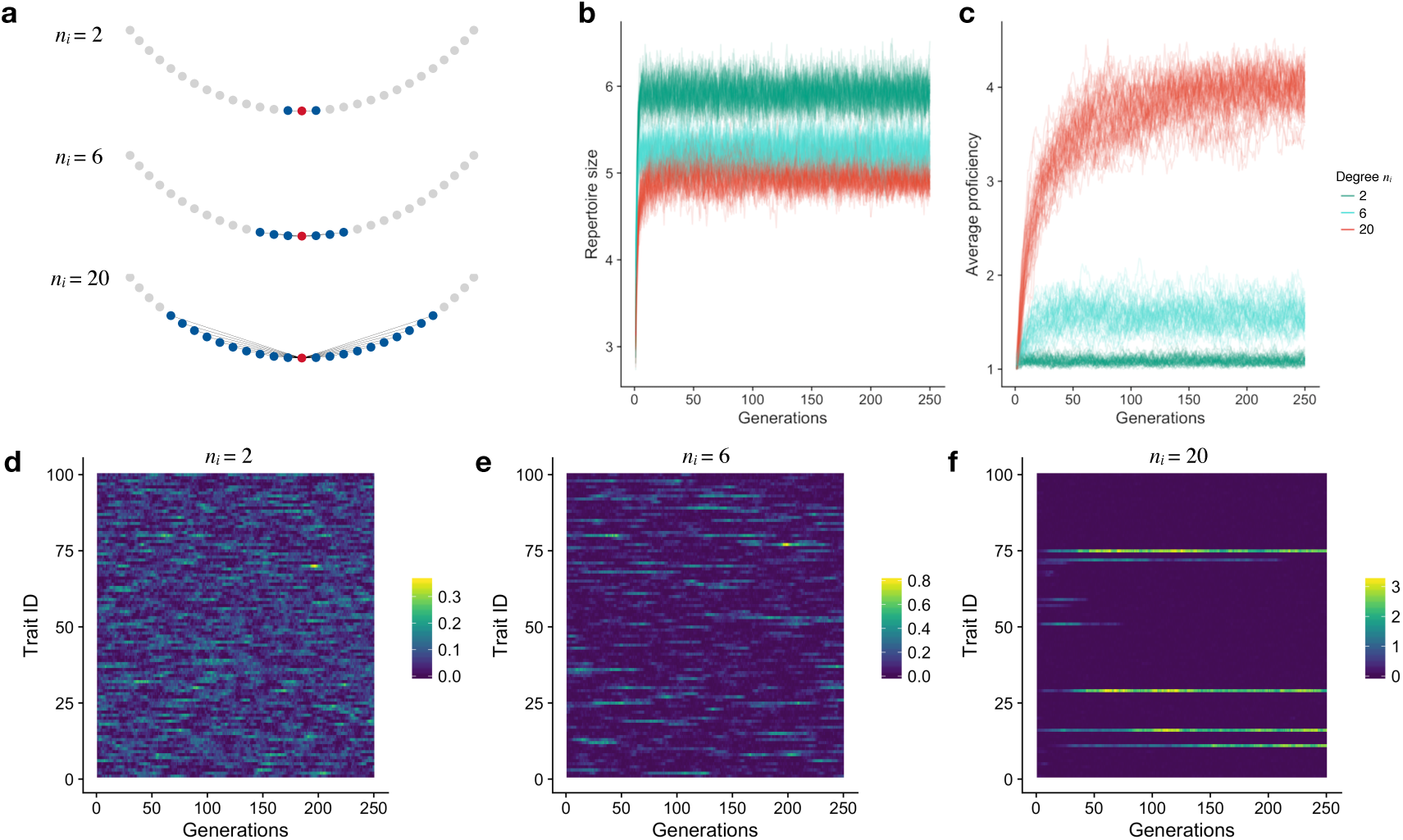
Increased connectivity leads to smaller trait diversity but higher trait proficiency. **a** Depending on the neighbourhood size *n_i_* a focal individual (red) is connected to 2, 6, or 20 neighbours (blue). **b** and **c**, Average repertoire size and average highest trait proficiency in populations with varying *n_i_*. Populations with larger *n_i_* (higher degree and shorter path length) have on average smaller repertoires and higher trait proficiency. **d-f**, Example record of average proficiency of all traits in populations with varying *n_i_*. Highly connected populations collectively possess fewer traits but are more proficient at them than sparsely connected ones.

In all subsequent simulations, we consider complex dynamic networks. This method mimics real-world networks and allows the dynamical formation of locally and globally clustered networks in response to different selective regimes^17^. In complex networks a new individual inherits two genes from its parent: *p_n_* (probability to form connections with the neighbours of the parent), and *p_r_* (probability to form connections with other individuals that are not connected to the parent). Mutation occurs with probability *μ* =1, whereby mutated values are drawn from a normal distribution centred around the parent’s value with standard deviations 0.1 and 0.01 for *p_n_* and *p_r_* respectively. We also ran simulations with lower mutation rates (*μ* = 0.01) and find that the results hold (ESM S5). While in the main text, we assume that connections can be formed at no costs, we also ran simulations where each connection incurs a cost, and find that costs can turn populations into generalists even if they are under specialist selection (ESM S6).

To determine the effect of topology on culture in *dynamic*, *complex networks*, we let networks dynamically rewire but keep *p_n_* and *p_r_* fixed throughout the simulation (Fig. 2). For both static, regular and dynamic, complex graphs, selection is neutral and reproduction is random. Thus, the spread of cultural traits is only affected by network topology.

**Figure 2:**
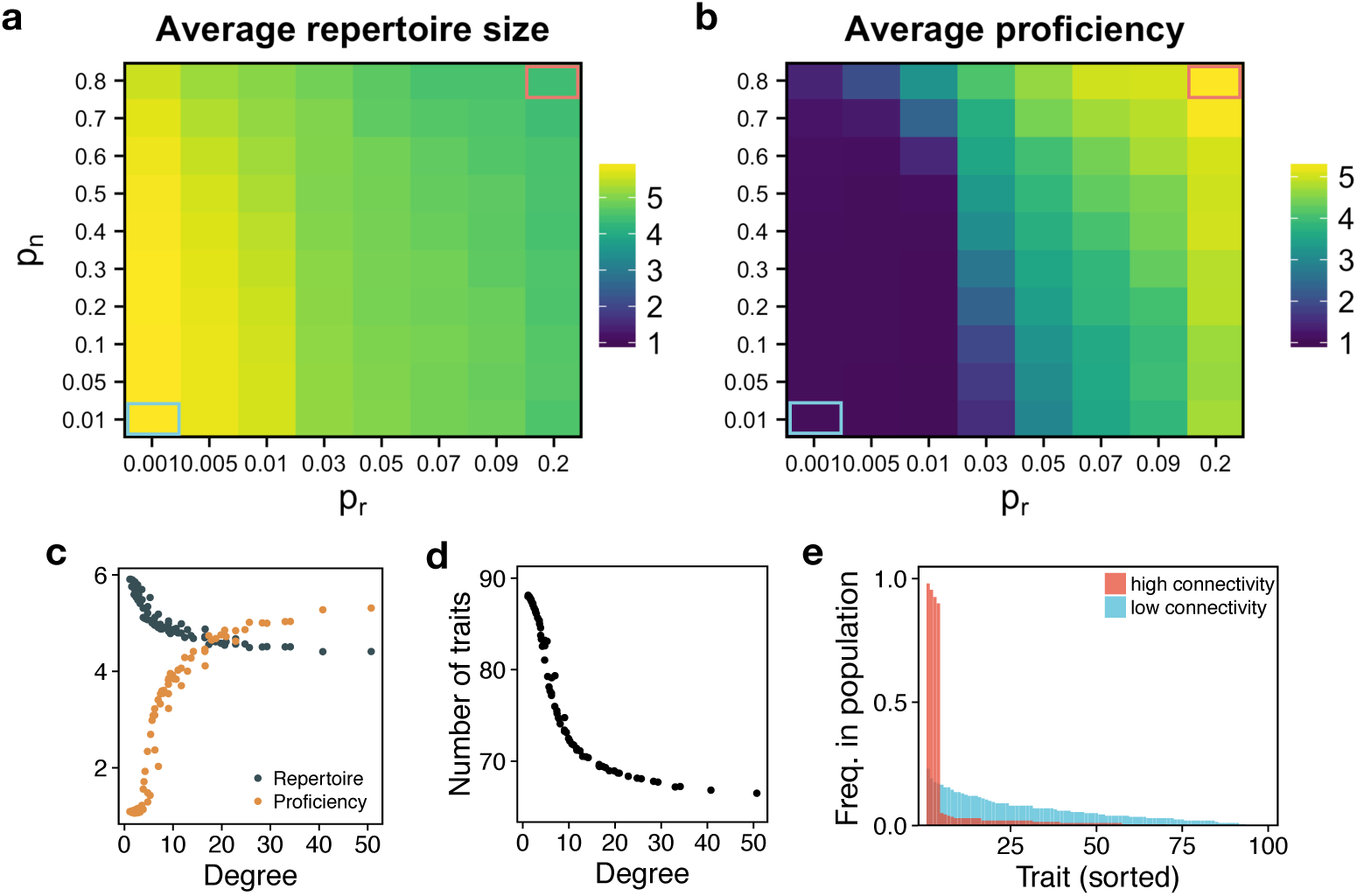
Effect of network topology onto cultural complexity. **a** and **c**, Sparsely connected populations (low *p_n_* and *p_r_*, low degree) have the largest individual repertoires, whereas well connected populations (high *p_n_* and *p*_r_, high degree) show highest trait proficiency, **b** and **c**. **d**, As average degree increases the total number of traits known to the population decreases. **e**, Moreover, the trait distribution for highly connected populations is skewed, such that a few traits are known to almost the entire population, whereas in sparsely connected populations traits are more evenly distributed, such that almost all available traits are known to different subsets of the population (compare lower left and upper right corner in **a** and **b**). All values represent population averages.

#### Co-evolution of network topology and culture

Next, we let both cultural knowledge and linking probabilities (*p_n_*, *p_r_*) evolve freely in response to the *generalists* or *specialists* environment (Fig. 3, see ESM S4 for time series).

**Figure 3:**
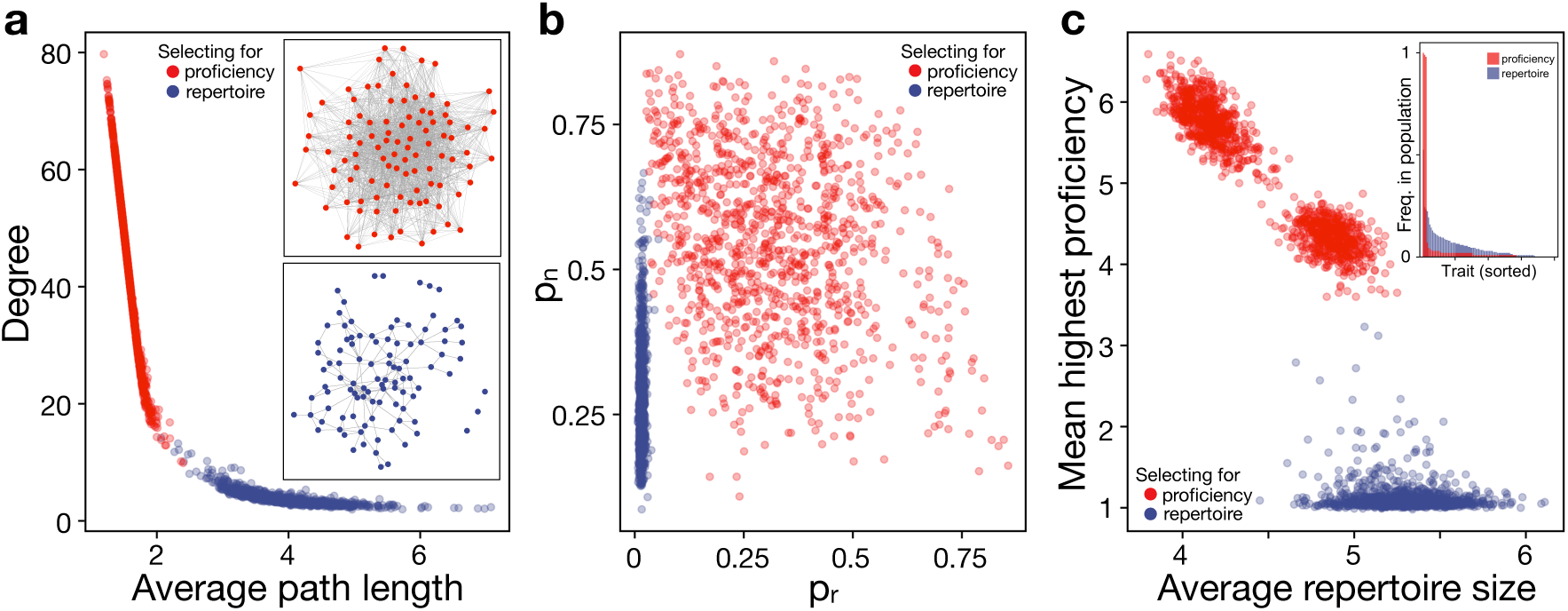
Specialist and generalist environments select for different network topologies. **a**, Selecting for trait proficiency increases degree and decreases average path length, leading to dense networks (example in upperinset), whereas, selecting for large repertoires decreases degree and increases path length, resulting in sparse networks (example in lower inset). **b**, While generalist environments strongly select against random connections (*p_r_*), there is little selection against either linking parameter in specialist environments. **c**, Selecting for generalists increases repertoire size while keeping proficiency at a minimum. Interestingly, selecting for specialists reduces repertoire size only slightly but strongly increased individual proficiency. Also, in simulations selecting for specialists results are distributed in two clusters. Here, populations differ in the number of traits they converged on. Populations with higher proficiency converged on three traits, whereas populations with lower proficiency converged on four traits (note, average repertoire sizes are one trait larger due to trait innovation). Under specialist selection trait distribution is highly skewed compared to generalist selection (inset).

#### Cultural response to enforced network degree

In nature, degree centrality might be more strongly affected by external factors than cultural selection. For example, sparse networks might be the result of high costs to form and maintain social connections. Networks might also be dense due to social norms or simply the spatial distribution of individuals. In both cases, this can lead to sub-optimal network densities. In this iteration, we investigate the trade-off between socially inherited and random connections when average degree is fixed. We run simulations where *p_n_* and *p_r_* are coupled as to achieve a pre-set degree centrality *k* (for calculations see ESM S3). A newborn still inherits *p_n_* from its parent, however, *p_r_* is based on a linear function with an inclination that depends on degree *k*. Should an individual form more than *k* connections, *k* − 1 connections are randomly chosen (one connection remains for the parent) and the rest discarded. We simulate populations with low, intermediate, and high connectivity (*k* ∈ 2, 6, 10) for both environments (Fig. 4).

**Figure 4:**
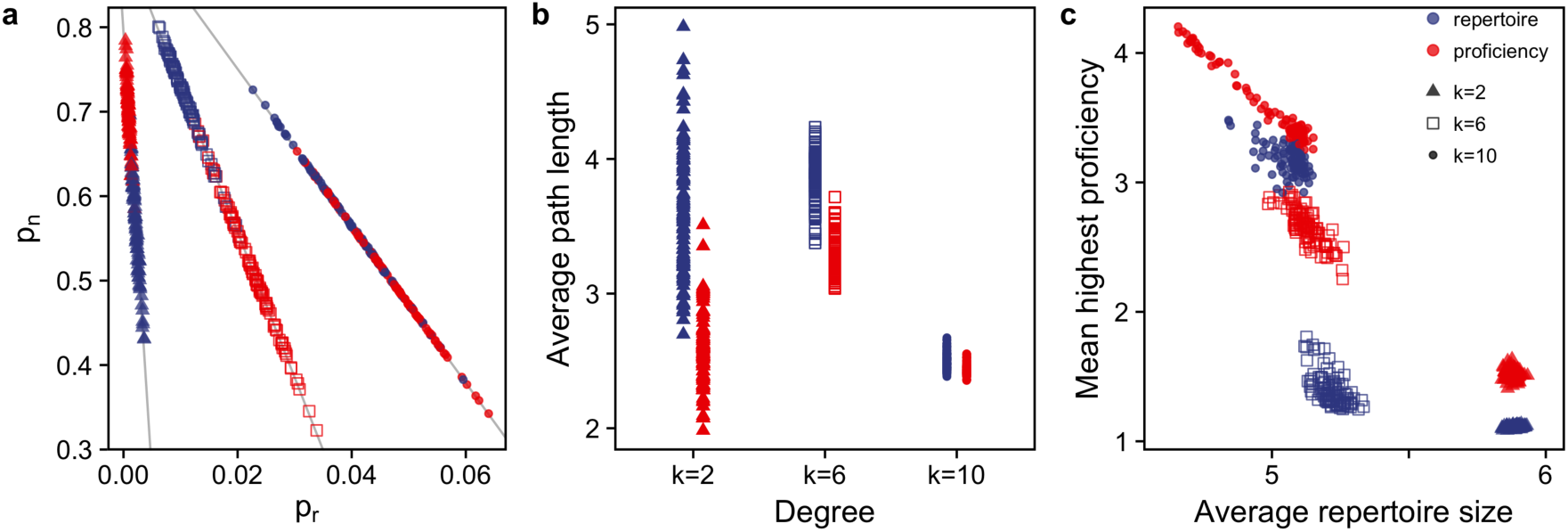
Trade-off between social inheritance and random connections depends on average connectivity. **a**, Specialists benefit from convergence of traits in their neighbourhood, which is achieved by increasing *p_n_* (at low connectivity) or increasing *p_r_* (at high connectivity). The opposite is the case for generalists, who try to avoid trait convergence in their neighbourhood. **b**, This leads on average to shorter path lengths in specialists and longer paths in generalists (lines indicate possible combinations of *p_n_* and *p_r_* given degree *k*). **c**, Populations are mainly made up of generalists at low connectivity and specialists at high connectivity. At intermediate connectivity, this is mediated by path length and clustering.

#### Cultural response to shifting environments

In a final set of simulations, we switch between both selective environments and observe the changes in topology and cultural repertoire (Fig. 5).

**Figure 5:**
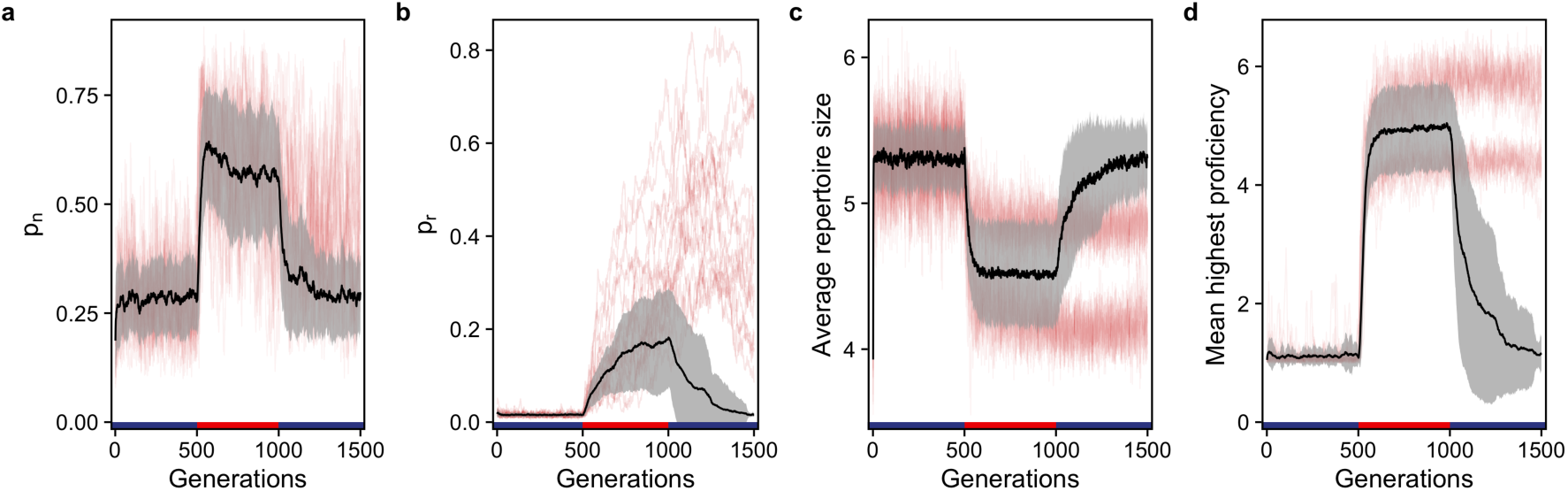
Populations more easily transition from generalists to specialists than the other way around. **a-d**, Trajectory of linking probabilities (*p_n_* and *p_r_*), and culture measures (repertoire size and proficiency), as selection regimes change from generalist (blue bars) to specialist (red bar) and back. Shown are mean (black line) and standard deviation (grey shading) for 100 repetitions. Some simulations do not return to larger repertoires and lower proficiency after the specialist phase (red lines). Here, although *p_n_* decreases after increasing in the specialist phase, **a**, *p_r_* keeps increasing even after selection is switched back to favouring generalists, **b**, see text for details.

### Parameters

If not stated otherwise, we run all simulations with *N* = 100 and *T* = 100 (see ESM S7 for additional values), for 1,000 generations (N death-birth events, with data being averaged over the last 200 generations) and 100 repetitions, with mutation rate *μ* = 1, innovation success rate *γ* = 0.01 and social learning success rate *σ* = 0.75 (see ESM S8 for additional values). Complex networks are initialised with *p_n_* = 0.1 and *p_r_* = 0.01. To compare differences in cultural knowledge between populations we record average repertoire size, and mean highest per individual trait proficiency. To compare networks, we record degree centrality, local clustering, and average path length.

## Results

### Network effects on cultural diversity

When we let culture evolve but keep the underlying social network fixed we find that average degree affects repertoire size and trait proficiency at the individual level (Fig. 1b-c), and trait diversity at the population level (Fig. 1d-f). Individuals in denser networks have higher trait proficiency and smaller repertoires, whereas those in more sparsely connected groups have larger trait repertoires but lower proficiency. Repertoire size and trait proficiency are negatively related due to the trade-off between spending learning turns on acquiring or improving traits. Also due to finite population sampling effects observing traits from more individuals is more likely to reinforce an existing skew in the trait distribution among neighbours. Consider a population where each individual possesses one out of 100 possible traits. The probability that two random neighbours have the same trait is 0.01. For six neighbours the probability that at least one trait appears multiple times grows to 0.14 and to 0.87 for 20 neighbours. As a result, we observe that dense networks have more skewed trait distributions and lower trait diversity than populations with sparse networks. In more connected groups proficiency is higher because the lower trait diversity increases the chance to repeatedly engage with the same trait, which increases proficiency (see eq. 2). In dense networks each learning phase acts as filtering process where rare traits will be learned less frequently, which further skews the distribution of traits on a population level leading to the loss of traits (Fig. 1f), but also a steady, accumulation of trait proficiency (Fig. 1b). In contrast, trait distribution is less skewed in sparsely connected networks. Individuals with low degree centrality are more likely to have neighbours with a variety of traits, which lets the observer build up a larger repertoire. However, because individuals rarely engage repeatedly with the same trait proficiency remains low. At the population-level, this keeps average trait diversity high but prohibits proficiency increase.

Interestingly, while average repertoire size plateaus very quickly average highest proficiency increases more slowly over the course of many generations (Fig. 1c). Because an observer’s trait proficiency cannot exceed the proficiency of the observed individuals through social learning alone, and given that innovations are rare, only populations with low trait diversity accumulate trait proficiency over time. Therefore, proficiency is higher in more connected populations.

### Dense networks have lower trait diversity and higher proficiency than sparse networks

We observe the same patterns when we apply the cultural dynamics to complex networks with dynamic rewiring but with fixed probabilities of linking *p_n_* and *p_r_* (Fig. 2). For high *p_n_* and *p_r_* (high degree and clustering, short average path length, see ESM S2) individuals have high proficiency and small repertoire sizes, whereas for small *p_n_* and *p_r_* (low degree and clustering, long average path length) individuals have larger repertoires but lower proficiency (Fig. 2c).

Repertoire size and trait proficiency are more affected by the rate of random linking (*p_r_*) as compared to the rate of inheriting connections (*p_n_*) because average path length is most strongly affected by *p_r_*. Longer average path length creates isolation by distance, which prevents the spread of a single set of traits throughout the entire network and so avoids population-wide coordination on a few traits. As with fixed networks, we observe strong differences in trait diversity between tightly and sparsely connected networks, with very few very common traits in highly connected populations contrasting a widespread and distributed knowledge of traits in sparsely connected populations (Fig. 2d,e).

### Specialists form dense, efficient networks, whereas generalists form sparse, inefficient networks

Next, we let linking probabilities *p_n_* and *p_r_* evolve in response to selection for specialist (favouring high trait proficiency) or generalist (favouring large repertoire size) knowledge. When selection favours specialist knowledge, populations evolve to form dense networks with high average degree and short average path length (Fig. 3a). Conversely, selecting for generalist knowledge yields sparse networks with low average degree and long average path length.

Interestingly, while in environments favouring specialization both linking probabilities appear to be under relatively weak selection, we find that there is strong selection against random connections when generalists are favoured (Fig. 3b). This may at first appear counter-intuitive, as random connections in a population with high trait diversity would allow individuals to be exposed to a completely different set of traits and as a consequence learn more different traits. However, we assume that successful social learning requires sufficient exposure to a trait (eq. 2). Therefore, generalists have to form neighbourhoods that trade-off trait homogeneity to increase trait exposure with trait diversity to increase chances to observe different traits.

Selecting for generalists or specialists has a marked effect on the culture in each environment (Fig. 3c). Selecting for generalists results in populations with larger trait repertoire but low proficiency, and selecting for specialists results in populations with fewer traits but higher proficiency. Although there is a 4- to 5-fold gap in proficiency between specialists and generalists, the difference in average individual repertoire size is relatively small, less than two traits on average. This reflects a reduction in overall learning in the high-diversity environment of generalists, where they do not get sufficient exposure to any single trait to attain high proficiency. In return, generalist populations as a whole carry many more traits at appreciable frequency (see inset in Fig. 3c), whereas specialists all learn the same few traits, resulting in a highly skewed trait distribution. This skewed trait distribution prohibits specialists from increasing their repertoire size, as all their neighbours converge to a small set of traits, whereas the broad trait distribution prohibits generalists from increasing their proficiency.

### Connectivity affects whether populations are generalists or specialists

The previous results show that generalists have a lower degree centrality than specialists. However, when degree centrality is externally dictated (high, intermediate, or low) individuals have to choose how many of their connections should be random links and how many should be socially inherited.

When degree centrality is small (*k* = 2), specialists are more likely to inherit connections than is the case for generalists (Fig. 4a). This is because specialists require a neighbourhood with low trait diversity, and given that they always connect to their parents it is more likely to find similar traits among their parent’s neighbours. For generalists, the opposite is the case. They are less likely to inherit connections and so increase the likelihood of learning from individuals with different trait sets. Therefore, average path length are longer in generalists than in specialists (Fig. 4b). In both cases we find populations that are made of small clusters. However, generalists have a wider variety of traits present in those clusters than is the case for specialists (Fig. S4a,d), leading to slightly higher proficiency for specialists (Fig. 4c and S5a,d).

When degree centrality is intermediate (*k* = 6) we observe the reversed pattern. Here, generalists are more likely to inherit connections than is the case for specialists. At this level of connectedness, specialists benefit from trait convergence, which allows an increased trait proficiency (Fig. 4c). Generalists avoid trait convergence in their neighbourhoods by increasing *p_n_* and formation of loosely connected clusters, which increases average path length (Fig. 4b).

At an even higher degree centrality (*k* = 10) generalists cannot avoid some convergence of traits (Fig. S4c), which leads to an overall increase in proficiency (Fig. 4c).

This shows that depending on the average degree a population will either mostly be made up of generalists (*k* = 2) or specialists (*k* = 10), while for intermediate degrees (*k* = 6) both states are possible and are mediated by average path length and clustering (Fig. 4c).

### When selection regimes change, specialists take longer to adapt than generalists

Switching between generalist and specialist environments reveals that the transition from sparsely connected populations with a broad distribution of traits occurs more readily than the transition from densely connected populations with highly specialised knowledge (Fig. 5). Subsequent to the switch to the specialist environment, average connectivity increases as *p_n_* and *p_r_* rise. This is followed by a decrease in average repertoire size and an increase in proficiency. Again, *p_r_* starts to drift as populations specialise on a few traits (Fig. 5b), leading to increasingly dense networks. When the environment changes again and favours generalists, not all populations return to a less connected state, but instead *p_r_* remains high (red lines in Fig. 5), and thus populations remain in a state of specialisation. This is due to an echo chamber like effect, where all individuals possess an almost identical set of traits and so a newborn will learn those common traits and improve its proficiency, leaving little to no learning attempts to acquire more traits. Novel traits are still innovated, but they are rarely copied by others. A change can only happen in small, isolated clusters. Here, individuals with new innovations can escape the conformity pressure from the rest of the population. Eventually, given sufficient time, populations will return to a more sparsely connected structure with higher trait diversity.

## Discussion

In this study, we combine cultural dynamics with evolving dynamic social networks to study how culturally mediated natural selection affects network structure. Our model highlights how selective effects of cumulative culture acquired from social connections determine both network structure and the diversity of cumulative culture at the individual and population level. We show that selection for generalist or specialist knowledge causes different network structures to emerge, with unexpected feedback onto population-level social and cultural dynamics. One might intuitively expect that selection for specialization might create internally clustered communities with sparse connections between them, whereas selection for generalists would favour more promiscuous connections to encounter a greater diversity of traits. In fact, we found the opposite effects: selection for generalists produces sparsely connected networks with few random links, whereas specialists increase their random linking to generate densely connected networks.

### Cultural-evolutionary feedbacks between network structure and cumulative culture

Our counter-intuitive results are driven by two factors, one at the individual level, the other at the network. At the individual level, we assume that successful social learning requires repeated engagement with the same trait, corresponding to the fact that learning in nature takes time and does not happen at the first contact with a novel trait^38–41^. Our learning model is therefore akin to complex contagion transmission of behaviour^18^, and differs from previous cultural learning models that largely relied on simple infection contagion^45;46^. A corollary of this learning model is that connecting with individuals who do not share many traits with each other depresses the overall rate of learning since no single trait is likely to be repeatedly observed. Therefore, overall learning rates are highest when connections share more traits, or in ecological terminology, when the beta diversity of traits within an individual’s neighbourhood is low. The second factor is the *cultural*-*evolutionary feedback* (similar to eco-evolutionary feedbacks) from network structure to the population-level cultural trait diversity. This feedback means that densely connected populations converge to the same few traits whereas sparsely connected ones display more network-level diversity through isolation-by-distance. This strong trade-off between population trait diversity and trait proficiency is mostly an emergent property. Although we also have (by necessity) a similar trade-off at the individual level, it is rather weak: specialization reduces individual repertoire sizes only slightly.

When selection favours specialists, we initially observe the evolution of high social inheritance and networks with partially connected clusters that specialise on specific traits. But the high internal connectivity in these clusters leads to a loss of local trait diversity. Due to their success in specialization, these clusters grow, creating a skewed trait distribution at the population-level where most individuals acquire the same few traits. As a result, even randomly formed links become likely to connect to an individual with similar traits to those connected through social inheritance. In turn, this reduces selection against random links, which increases average connectivity, and further decreases trait diversity and skews trait distribution. The eventual outcome of this process is that almost all individuals will converge to an almost identical set of traits and therefore the number and type of links become essentially neutral.

Conversely, selecting for generalists favours the ability to learn rare traits (perhaps individually innovated by a given connection). But this is only possible when rare traits are relatively common within the local neighbourhood of a focal individual, which favours low connectivity (as it decreases the total number of traits amongst the connections of a focal individual). This puts pressure on overall connectivity to decrease, and through isolation-by-distance, increases the overall cultural diversity of the population. One might think that the increased cultural diversity might “tempt” individuals to make more random connections to access new rare traits (which would undermine trait diversity at the population level), but this temptation is counteracted by the fact that being connected to individuals with too much diversity in traits depresses the overall learning rate.

### Exploration and coordination on social networks

Our results inform several important questions in the study of cultural evolution. For instance, much work has focused on how network structure affects the emergence and spread of social conventions and norms. While our model does not have direct social effects on payoff (i.e., the payoff of an individual depends only on its own trait), it nonetheless captures the same cultural-evolutionary dynamics underlying social norms and social information. Our results parallel recent experimental evidence showing that social conventions can emerge spontaneously and groups converge quickly on norms if the underlying social network is well connected^47^. Another important question is how social network structure affects information aggregation. Social influence can bias individual estimates^48^ but whether such biases improve or undermine the “wisdom of the crowd” depends on the underlying social network. A recent study found that decentralised groups became more accurate over time, whereas centralised groups, where central individuals have a disproportionally large effect on the collective estimation process, were more likely to increase in error^49^. Further, a recent lab experiment showed that a group’s performance in finding solutions to a complex problem can initially profit from dense information networks. However, the fast dissemination of successful solutions decreased exploration of the solution space and made well-connected groups more likely to settle for suboptimal local maxima^27^. Our theoretical results are consistent with these findings, and further imply that selection for proficiency can in the long-term lock populations into well-connected networks which on one hand might increase their ability to coordinate on a set of conventions, but on the other hand diminish their capacity to explore or incorporate new information.

A related phenomenon is the spread of information and opinions on social media. Information that is shared in social media spreads in a complex contagion manner^50^, where the likelihood an individual will spread information increases monotonically with exposure^51^. Previous research has connected homophily of connections (individuals tend to be connected with like-minded individuals) to the emergence of “echo chambers” where the same information gets shared over a network^52^. Here, we show that the cultural-evolutionary feedbacks with complex contagion-like learning can produce convergence on the same few traits in well connected networks even without explicit homophily.

An interesting area for studying cultural-evolutionary feedbacks is how scientific fields self-organize and explore different questions and methods. As research fields grow, networked communities of researchers emerge and the shape of these networks affects the spread of questions, study designs, and analytic methods. A recent study showed that centralized research communities were less likely to produce replicable results^53^. In contrast, decentralised communities (connecting researchers from independent groups) relied on more diverse methodologies and generated more robust and replicable results^53^. These results are consistent with our model which shows that low path length networks (i.e., centralized) tend to converge to a few traits. Studying the feedback between how evolving scientific networks and specialization into different scientific “traits” will likely illuminate how scientific discovery proceeds.

### Population size, networks, and cultural complexity

One of the ongoing debates in cultural evolution is the role of population size^4;5;54^ versus connectivity and mobility^7;55^ in determining the cultural diversity and complexity. We find that larger populations maintain more traits (see ESM S7), but more connected groups achieve higher proficiency at the cost of overall trait diversity. Interestingly, we find that cultural-evolutionary feedbacks cause the transition between the generalist, highly diverse and specialist, highly proficient states to be highly non-linear. At low connectivity proficiency is low and trait diversity is high. When the network reaches a critical density (here, at an average degree of about 5), we find a sudden increase in trait proficiency. Due to the increased number of individuals learning and innovating along the same traits, trait proficiency increases. These dynamics can be interpreted as an increase in the ‘effective cultural population size’ (for a few traits) which has been suggested as the main driver of the transition between Middle and Upper Palaeolithic, marked by an increase technological complexity but also by increased inter-connectedness between groups^56^. Consistent with our results, archaeological analysis suggests that increased inter-group connectedness lead to decreased technological volatility^57^.

Our simulations also show an upper limit for the number of traits an individual can acquire, and by extension, an upper limit of traits a population can carry. However, as part of cumulative cultural evolution traits not only become more complex but also increase in number^58^. To allow culture to continuously expand in our model, traits could directly affect learning, for example, by making it easier to learn a trait (less costly, see ref. 59), or by directly affecting demography, for example, by increasing carrying capacity^56;60^.

## Conclusion

We find that network topology not only affects the diversity and accumulation of cultural knowledge but is itself shaped by cultural selection, due to a continuous eco-evolutionary feedback between social structure and culture. As we have shown, it is important to model individual-level interactions to understand this feedback. Recent technological advances allow us to gather detailed individual data^61^. Further empirical research, especially long-term studies, will help to clarify the extent of cultural selection on human social networks, and thus shed light on the origins of cumulative culture in our ancestors.

## Code availability

All code used in this paper is available at (URL added before submission).

## Author contributions

E.A. secured funding. M.S. and E.A. contributed to the study design, M.S. conducted analyses,
M.S. wrote the initial draft of the manuscript and all authors contributed to revisions.

## Funding

M.S. was supported by a grant to Erol Akçay from the Army Research Office (W911NF-12-R-0012-03).

## Competing interests

The authors declare no competing interests.

## Additional information

Supplementary information is available for this paper at *URL*.

